# Biochemical and kinetic properties of a Type III restriction-modification enzyme Mbo45V from the host-adapted pathogen *Mycoplasma bovis*

**DOI:** 10.64898/2026.05.01.722158

**Authors:** Ishtiyaq Ahmed, Akhilesh Pratap Singh, Om Prakash Chauhan, Karishma Bhagat, Aathira Gopinath, Kayarat Saikrishnan

## Abstract

Type III restriction-modification (RM) enzymes are prominent bacterial defense against bacteriophage and invading foreign DNA that also modulate the host’s epigenetic landscape. Genome analysis of the host-adapted *Mycoplasma bovis* PG45 that has a very small genome revealed a Type III RM locus comprising one res and three mod genes. We characterized Mbo45V, a representative enzyme encoded by this locus. The enzyme forms a heterotrimeric complex consisting of two Mod subunits and one Res subunit. Mbo45V recognizes the asymmetric sequence 5′-YAATC-3′ (Y = T/C) and cleaves DNA having at least two head-to-head oriented sites ∼26–28 bp away from the recognition site. Methylation of the second adenine of the target site using cofactor S-adenosylmethionine (SAM) protects DNA from restriction, while the SAM analogue sinefungin enhances DNA binding and cleavage. Kinetic studies reveal that Mbo45V exhibits relatively weak DNA binding affinity and an unusually high K_m_ for SAM, indicating low cofactor affinity compared to prototypical enzymes such as EcoP15I. ATPase activity is strongly stimulated by cognate DNA and is inhibited upon methylation of the substrate, suggesting a regulatory interplay between methylation and restriction functions. Comparative analysis indicates that, although Mbo45V shares core mechanistic features with prototypes from *Escherichia coli*, its kinetic parameters are distinct. These differences likely reflect adaptation to the stable intracellular environment of *M. bovis*, in contrast to the fluctuating conditions encountered by the enteric bacteria.

## Introduction

*Mycoplasma* is an important genus of bacteria, which is the causative agent of many diseases affecting humans and animals. For example, *Mycoplasma bovis* causes bovine pneumonia, mastitis, and arthritis in cattle and is a major concern for cattle-based industry and livelihood worldwide. Organisms belonging to the genus *Mycoplasma* lacks cell wall, have a minimal genome, a reduced coding capacity, and a limited number of metabolic pathways^1^. The small genome size has led to the view that most retained genes are essential for survival in their natural habitats. Most species are obligate parasites that thrive within their hosts. Given their compact genomes encoding far fewer proteins than other bacteria, it is notable that they harbor multiple genes for restriction–modification (RM) enzymes^2,3^, emphasizing the importance of these enzymes for pathogen survival in the host.

RM enzymes are a bacterial defense that prevent entry of foreign DNA into host by degrading the DNA. The RM enzymes can protect bacteria from infectious bacteriophages or prevent acquisition of exogenous DNA from the environment, including antibiotic resistance genes and pathogenicity islands^4^. They have two components, a methyltransferase (MTase) that methylates (modifies) the host genome at specific recognition sites and an endonuclease that cleaves DNA having unmodified recognition sites. Based on their cofactor requirement, subunit assembly and the mode of endonucleolytic cleavage, RM enzymes are classified primarily into four types – Type I, Type ISP, Type II and Type III^5^.

Type III RM enzymes, the focus of this study, consist of Mod and Res subunits^6^,^7^. The Mod subunit contains the core MTase domain and Target Recognition Domain (TRD), which facilitates recognition of the target sequence. The Res subunit is made of a nuclease domain coupled to an ATPase belonging to the Superfamily 2 of helicases. Based on the studies of the prototypical EcoP1 and EcoP15I, we know that Type III RM enzymes efficiently cleave DNA with at least two recognition sites in an inverted orientation separated by thousands of base pairs^6^, with cleavage occurring close to one of the two recognition sites^8^. Both the enzymes can also cleave single-site substrates, albeit with lower efficiency^9^.

Type III RM enzymes are known to perform another important function – to alter the host’s gene expression. Alteration in expression is achieved through the enzyme’s DNA methylation activity^10^. Additionally, in host-adapted pathogens, such as *Neisseria, Haemophilus, Moraxella, Mycoplasma*, etc., the Type III RM enzymes are subjected to phase variation. Phase variation is the high-frequency reversible on-off switching of a gene ^11^(10). Phase variation of the Type III RM enzymes occur due to high frequency indel formation in the small sequence repeats (SSR) present in the open reading frame (ORF) of the *mod* gene^12^.

In host-adapted pathogens, phase variation is a means of introducing phenotypic diversity and increasing pathogenicity, which facilitates colonization of the host, adaptation to the host environment and evasion of the host immune system^13^. RM systems, such as the Type I, Type ISP, Type IIL and Type III systems, in which the MTase is integral to DNA recognition and the endonuclease activity can function as phase variable systems, since turning off the MTase or changing its specificity simultaneously also alters the endonuclease activity^13^.

As the MTase activity of Type III RM enzymes regulate the expression of other genes, phase-variable *mod* genes can likewise alter the expression of these genes too. The phase-variable RM systems are proposed to (i) confer phase-variable resistance to phage infection; (ii) facilitate acquisition of potentially useful foreign DNA by the temporary loss of action of the RM enzymes^14,15^; (iii) promote degradation of the genomic DNA of the host organism with consequences to the fitness level of the population^16^; and (iv) regulate gene expression via differential methylation of the genome^17,18,19^.

Among Type III RM enzymes, only the prototypical and closely related EcoP1 and EcoP15I, encoded by prophages of *Escherichia coli*, have been biochemically and biophysically characterized in detail. Consequently, information on the biochemical and enzymatic properties of Type III RM enzymes from other organisms, particularly the host-adapted pathogens, is limited. Unlike *E. coli*, which is a facultative anaerobe that can live in diverse environments, including within or outside a host under fluctuating nutrient conditions, *M. bovis* resides always within the host in a nutrient-rich milieu.

Previously, Type III RM enzymes from have been identified in *Mycoplasma pulmonis* ^20^ and *Mycoplasma mycoides* subsp. *capri*^21^. Here, we describe the identification and comprehensive biochemical characterization of a potentially phase-variable Type III RM enzyme from *M. bovis*, designated Mbo45V. We report its DNA target sequence and cleavage mechanism, present detailed kinetic analyses of its activities, and compare its properties with those of EcoP15I and EcoPI from *E. coli*. The comparison reveals that the kinetic behaviour of Mbo45V differs substantially from that of well-characterized β-class methyltransferases. This study highlights both shared and distinct features of Type III RM enzymes from two phylogenetically and ecologically distinct bacteria—the obligate parasite *M. bovis* and *E. coli*.

## Materials and Methods

### DNA

All the oligonucleotides were purchased from Sigma-Aldrich or Integrated DNA Technologies. A 235 bp DNA substrate containing two recognition sites for Mbo45V oriented head-to-head was generated by PCR from a plasmid using primers For_Pri_235 and Rev_Pri_235 (Supplementary Table 1). The PCR amplified DNA substrate was purified using Qiagen PCR purification kit, and the concentration was determined using Thermo Scientific™NanoDrop™spectrophotometer. A 650 bp DNA containing a single recognition site was used to generate two-site DNA substrates by site-directed mutagenesis using the primers (i) For_Pri_650_1 in combination with For_Pri_650 and Rev_Pri_650 for head-to-head sites; (ii) Rev_Pri_650_2 for head-to-tail sites; (iii) For_Pri_650_3 for tail-to-tail sites. See Supplementary Table 1 for primer details.

### Isolation and cloning of Mbo45V

We amplified the *mod* and *res* genes as a single operon and as individual genes from the genomic DNA of *M. bovis* Donetta PG 45 obtained from DSMZ (Catalogue no. 22781). *M. bovis* uses TGA as the codon for tryptophan, which is a stop codon in *Escherichia coli*. Heterologous expression in *E. coli* of the *res* and *mod* genes from *M. bovis* required the mutation of all the thirteen TGA tryptophan codons to TGG to express the full-length protein. Also, the SSR was deleted by overlap PCR without affecting the ORF. The deletion of the SSR removed the Arg-Glu amino acid repeat. All the primers used for the mutagenesis are listed in Supplementary Table 2. After this, *mod* and *res* were cloned into two separate high expression vectors, respectively.

*mod* was cloned into pRSF using For_Mod_RSF and Rev_Mod_RSF (Supplementary Table 2). The amplified gene was double digested using the restriction enzymes NcoI and XhoI and ligated into pRSF vector. The recombinant vector pRSF*mod* was confirmed by DNA sequencing. Similarly, the *res* gene with all internal TGA codons changed to TGG was PCR amplified using For_Res_His and Rev_Res_His (see Supplementary Table 2 for primer details). The amplified *res* gene was double digested with NdeI and BamHI restriction enzyme and ligated into pHIS vector.

### Purification of Mbo45V Mod

*E. coli* BL21(DE3) cells were transformed with recombinant vector pRSF*mod* having kanamycin as a selection marker. A 4 L culture of 1x LB was grown at 37°C until the OD_600_ of the culture reached 0.5. The culture was shifted to 18°C and induced at an OD_600_ of 0.6 with 1 mM IPTG. The culture was further grown for 12 h after which the culture was harvested and pelleted down. The culture was resuspended in 200 mL lysis buffer (50 mM Tris-Cl pH 8, 500 mM NaCl, 10% glycerol, 10 mM MgCl_2_, and 1 mM DTT). The resuspended culture was lysed by sonication in a 3 mins cycle with 1 sec on and 3 seconds off. To ensure complete lysis of the cell, the sonication cycle was repeated two times. Also, to prevent heating of the sample during sonication, the entire process was carried out at 4°C. The lysate was spun at 100,000 g at 4°C for 1 h using Optima XE ultracentrifuge (Beckman Coulter). The clarified supernatant was loaded on to the 5 mL Ni-NTA column pre-equilibrated with Buffer A (50 mM Tris-HCl pH 8, 500 mM NaCl, 15 mM imidazole). The protein was eluted from the Ni-NTA column using a stepwise gradient of higher concentration of imidazole Buffer B (50 mM Tris-HCl pH 8, 500 mM NaCl, 500 mM imidazole). The eluted fractions containing protein were pooled and dialyzed against buffer B-50 (50 mM Tris-HCl pH 8, 50 mM NaCl, 1 mM DTT) for 8 h at 4°C. After dialysis, the protein was centrifuged using Avanti J-26XP (Beckman Coulter Life Sciences) for 30 minutes at 17418 rcf and 4°C to remove any aggregates and particles. The protein was loaded onto a Mono Q 10/100 GL column (GE Healthcare) pre-equilibrated with B-50. The protein was eluted from Mono Q 10/100 GL by a linear gradient from 50 to 500 mM NaCl in buffer B. The fractions containing purified Mbo45V Mod protein were pooled together and concentrated using Viva spin column (MWCO 30 kDa, GE healthcare). The last step of purification SEC using a Superdex 200 10/300 (GE Healthcare) column was carried out. The fractions from SEC were pooled together and concentrated using Viva spin column (MWCO 30 kDa, GE healthcare) and stored at -80°C until further use.

### Purification of Mbo45V

Recombinant vectors pRSF*mod* and pHIS*res* were co-transformed into *E. coli* BL21(DE3) cells with both ampicillin (100 µg/ml) and kanamycin (50 µg/ml) as a selection marker. 6 L of 1x LB was grown at 37°C with constant shaking at 250 rpm. The culture was grown until an OD_600_ of 0.6 was reached after which the culture was shifted to 18°C and induced with 2 g/L of L-arabinose and 1 mM IPTG. The culture was harvested after 12 h, pelleted down and resuspended into 200 ml of lysis buffer [50 mM Tris-Cl pH 8, 500 mM NaCl, 10% glycerol, 10 mM MgCl_2_, and 1 mM DTT]. The resuspended culture was lysed by sonication at 4°C, with 1 sec ON and 3sec OFF, and the sonication cycle was repeated two times. The lysate was spun at 37000 rpm at 4°C for 1 h using Optima XE ultracentrifuge (Beckman Coulter). The resulting supernatant was loaded onto a 5 ml Ni-NTA column pre-equilibrated with Buffer A [50 mM Tris-HCl pH 8, 500 mM NaCl, 10 mM imidazole]. To eliminate the impurities bound non-specifically to the column, washes of 4 column volumes (CV) of 3% and 8% Buffer B [50 mM Tris-HCl pH 8, 500 mM NaCl, 500 mM imidazole] was performed. The protein was eluted with an increasing concentration of imidazole in a stepwise gradient using a mix of Buffer A and B. The eluted fractions containing Mbo45V protein were pooled together and dialyzed against B-50 (50 mM Tris-HCl pH 8, 50 mM NaCl, 1 mM DTT) for 6 hours at 4°C. The dialyzed protein was centrifuged for 20 minutes using Avanti J-26XP (Beckman Coulter Life Sciences) at 4°C. The dialyzed protein was further purified using a Mono Q 10/100 GL column (GE Healthcare) pre-equilibrated with B-50 buffer. Mbo45V was eluted from the column using a linear gradient of NaCl (50 mM to 500 mM). Mbo45V eluted from Mono Q 10/100 GL column at ∼280 mM NaCl. Size exclusion chromatography (SEC) using Superdex 200 10/300 (GE Healthcare) was carried out as the last step to check the homogeneity of Mbo45V. Fractions containing pure protein were pooled, concentrated using centrifuge concentrator Vivaspin column (MWCO 30kDa, GE healthcare) and stored at -80°C until further use. Activity test of the protein was not carried out during the steps of purification. Only the final purified sample was used for all the subsequent enzymatic assays.

### Size-exclusion chromatography with multi-angle light scattering

Size-exclusion chromatography, combined with multi-angle light scattering (SEC-MALS) was used to determine the molecular weight of RM accurately. Mbo45V at 2 mg/ml was loaded on Superdex 200 10/300 (GE Healthcare) at a flow rate of 0.3 ml/min. The column was connected to a light scattering diode array and differential refractive index detector (Wyatt Technology UK Ltd). Before starting the experiment, BSA was run as a standard to check the reproducibility of the results. The chromatographs were analyzed using Graph Pad software.

### DNA Cleavage assay

DNA cleavage assay was carried out in a cleavage buffer [10 mM Tris-acetate pH 8, 10 mM potassium acetate, 10 mM magnesium acetate, 1 mM DTT] at 37°C. Protein and DNA were mixed in the presence of 20 µM SNF, and the reaction was started by the addition of 4 mM ATP. The reaction was further carried out for 45 minutes and stopped by the addition of 0.5X volume of STOP buffer (10 mM Tris-HCl pH 8, 100 mM EDTA, 20% v/v glycerol, 0.025% SDS, 0.03% bromophenol blue). The samples were heated at 65°C for 15 minutes to inactivate the enzyme and then loaded on to a 12% native PAGE gel.

### Identification of the Mbo45V cleavage loci

To locate the DNA cleavage position by Mbo45V, a 1 kb DNA substrate (50 nM) containing two recognition sites of Mbo45V in head-to-head orientation was treated with Mbo45V (300 nM) in cleavage buffer. The reaction was started by adding 4 mM ATP and carried out for 1 h at 37°C. The cleaved products were isolated using agarose gel, purified using the QIAGEN gel extraction kit, and analyzed by run-off sequencing.

### Electrophoretic Mobility Sift Assay (EMSA)

DNA binding ability of Protein to a 30 bp long DNA (Mbo45V_30mer) studied using Electrophoretic mobility shift assay (EMSA). Varying concentrations of the protein (0.25 to 2 μM) were incubated with 250 nM DNA in a binding buffer containing 10 mM Tris-acetate pH 8, 10 mM potassium acetate, 10 mM magnesium acetate and 1 mM DTT at 37°C for 20 mins. The reaction was stopped by the addition of 0.4 X volume of stop buffer containing 10 mM Tris-acetate pH 8, 10% v/v glycerol, 20% w/v sucrose. All the samples were loaded on to a 5% native gel, which was run at 4°C in 1X Tris-Borate-EDTA (TBE) buffer with additional 5% glycerol in the running buffer.

### Fluorescence anisotropy

Fluorescence anisotropy measures the polarization of an emitted beam of light from a fluorescent molecule that has been excited with a polarised beam of light. Fluorophore with high rotation diffusion will have low anisotropy. Rotational diffusion of fluorophore inversely depends on the size of the fluorophore complex, so anisotropy is directly dependent on the size of the Fluorophore complex^23^. 5’-FAM labelled dsDNA ([6FAM] TGGCTT CAGCAG TAATCC GCAGAT ACCAAA ACTGTCCG) with cofactor and varying protein concentrations are mixed in reaction buffer. 200ul reaction mixture is used in the cuvette to record fluorescence anisotropy by Horiba Fluromax4. Anisotropy values are plotted against protein concentration. The plot was fit for a single site-specific binding equation (y (anisotropy) = B_max_[P] / k_d_+[P]) to get the dissociation constant (k_d_).

### DNA methylation assay

300 nM of Protein were incubated with 50 nM of 235 bp DNA substrate containing two recognition sites in head-to-head orientation at 37°C in the presence of 200 µM AdoMet for 1 h. The methylated DNA was purified, and the extent of DNA methylation was confirmed by carrying out DNA cleavage assay with Mbo45V.

### DNA Methylation kinetics

The MTase-Glo Methyltransferase Assay kit from Promega is used to measure the DNA methylation activity by enzymes. In this kit enzymes, convert SAH into ATP, which is <u>subsequently</u> utilized by luciferase to produce luminescence, so luminescence generated in the reaction is directly proportional to the presence of SAH^24^. To calculate the amount of SAH produced in the reaction. SAH standard is performed with each set of experiments. StD SAH concentration plotted against respective luminescence and fit against equation y = mx + C. Std slope value is used further to determine the amount of SAH produced in the reaction. We have incubated the 300 nM protein with varying concentrations of substrate (DNA or SAM) while keeping other substrates at saturated concentrations (DNA 3 mM or SAM 500 mM) for 1hrs. The rate of reaction is determined and plotted against the concentration of substrate. To determine the kinetic parameter plot was fit in equation V= Vmax[S]/ KM +[S].

### ATPase assay

ATP hydrolysis by protein was measured using a NADH coupled enzyme assay. The assay is based on ATP hydrolysis reaction, which is coupled to the oxidation reaction of NADH by pyruvate kinase (PK) and lactate dehydrogenase (LDH). Here, we measured decrease in NADH absorbance at 340 nm in a 96-well plate using Varioskan Flash (Thermo Scientific). The decrease in the NADH absorbance is proportional to the rate of ATP hydrolysis^25^. NADH, ATP and the enzymes PK and LDH were purchased from Sigma. A 69 bp long DNA having one recognition site of Mbo45V enzyme was used as a specific substrate, and the same length of DNA with no recognition sequence was used as non-specific DNA. The reaction mix had 600/300 nM of protein and 1 µM of DNA. To start the reaction 1 mM ATP was added, reaction was mixed and incubated for 30 s. Absorbance at 340 nm was recorded at an interval of 10 s for 360 s at 25^°^C. ADP standard was performed for each set of experiments. The concentration of ATP hydrolyzed was calculated at each time interval using a line equation Y= mX+ C (where Y= absorbance, m = slope, X = concentration of ADP produced or ATP hydrolyzed and C = intercept on Y axis) obtained from standard plot with different ADP concentrations (100 – 600 µM). To determine kinetic parameters 300 nM protein, 1mM DNA and varying concentration of ATP (0.001-1 mM) is used. Rate of ATP hydrolysis is plotted against the concentration of ATP. Plot was fitted on Equation V=V_max_[ATP]/K_M_+[ATP] to get the V_max_ and K_M_ for ATP hydrolysis.

## Results

### Organization of the genes encoding a Type III RM enzyme in M. bovis

Genome analysis of *M. bovis* Donetta PG 45 strain revealed a cluster of a *res* and three *mod* genes - *mod1, mod2* and *mod3* (locus Mbo45ORF167P in the REBASE database) (Figure 1A). A similar cluster of *res* and *mod* genes was found in the genome of *Mycoplasma pulmonis* (Dybvig et al., 2006). In *M. bovis* genome, the end of the *mod1* gene overlapped with the first 14 bp of the *res* gene. The overlap of *mod1* and *res genes* was similar to the organization of the prototypical EcoP1 and EcoP15I *mod* and *res* genes. Mod1 shared an amino acid sequence identity of 70.2% and 68.7% with Mod2 and Mod3, while Mod2 was 70.7% identical to Mod3.

**Figure 1.**
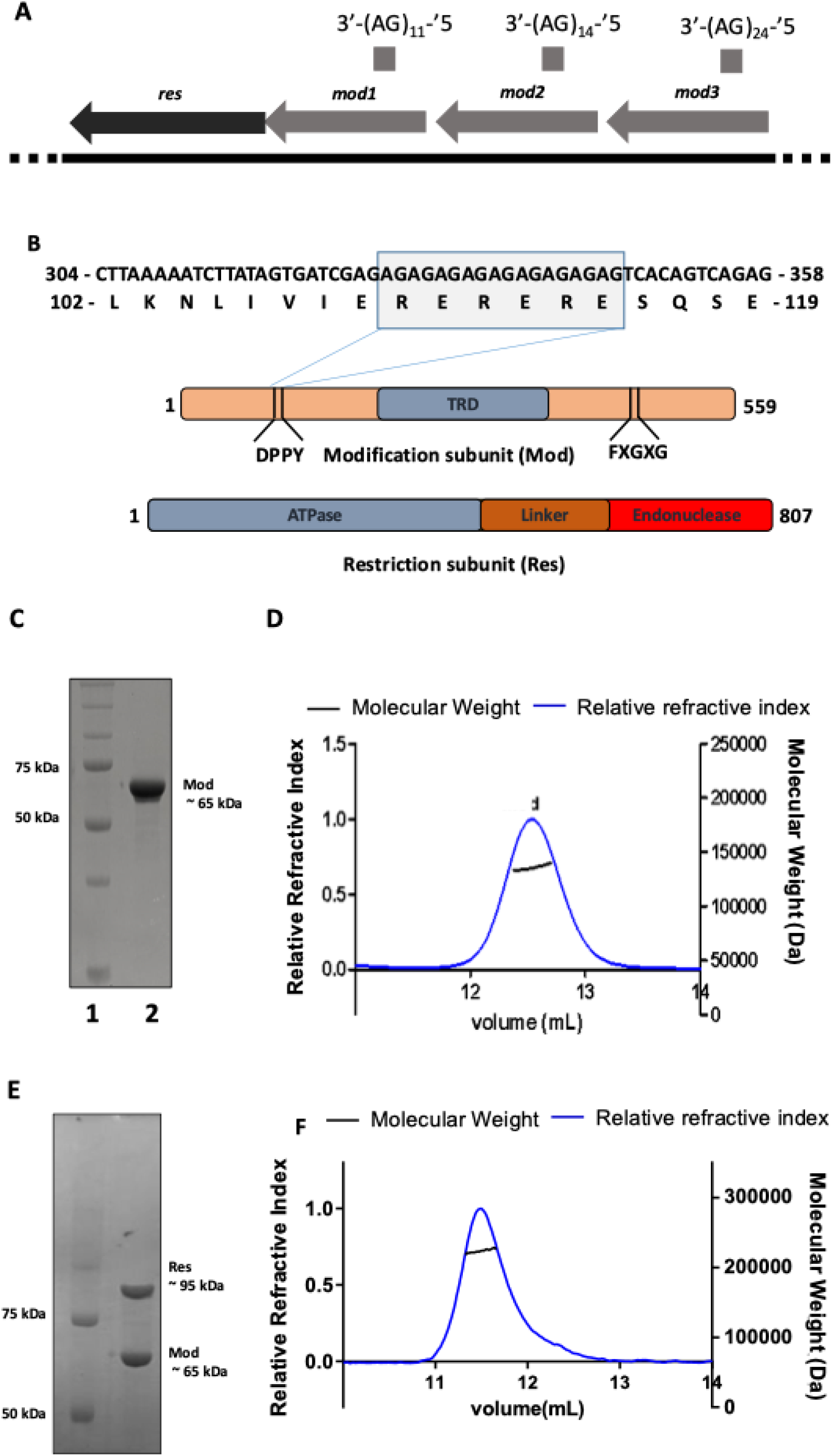
Domain organization and oligomeric state of MboIII. (A) Gene arrangement in operon *mbo45orf167P*, contain 3 *mod* and 1 *res* gene, mod1 had 11 AG SSR repeat, *mod2* had 14 AG SSR repeat and *mod3* had 24 AG repeat. (B) Domain organization of the Mod subunit illustrating the position of motif I (FXGXG), motif IV (DPPY), and the TRD. Domain organization of the Res subunit illustrating the position of the ATPase, the linker and the endonuclease domain. (C) SDS-PAGE analysis of purified Mod. Lane 1 is protein ladder, lane 2 is purified Mod subunit (M.Mbo45V) subsequent to SEC. (D) SEC-MALS elution profile of M.Mbo45V. Mod eluted as a single peak with a molecular weight of 131 kDa indicating that it is a homodimer. The solid line on left (Y-axis) represents the normalized light scattering while the dotted line on the right (Y-axis) represents the calculated molecular weight. (E) SDS-PAGE analysis of purified Mbo45V. Lane 1 is a protein ladder, lane 2 is the purified Mbo45V. (F) SEC-MALS elution profile of Mbo45V. Mbo45V eluted as a single peak with a molecular weight of 225 kDa indicating that it is a heterotrimer.

We amplified the *mod1* and *res* genes together as a single operon, and separately as individual genes from the genomic DNA. The 65.6 kDa protein product encoded by *mod1* had conserved motifs arranged as in a β-class of N^6^-adenine MTase (Figure 1B). *mod1* amplified from the genomic DNA had a SSR of 11 AG repeats, which would result in premature termination of translation. To translate the ORF, we generated two constructs – one having 9 AG repeats (*mod1*^9AG^) and the other having 0 repeats (*mod1*^0AG^) (Figure 1B). An amino acid sequence analysis of the 94.7 kDa protein encoded by *res* showed that it has all the motifs characteristic of an SF2 helicase at the N-terminus and an endonuclease domain at the C-terminus (Figure 1B).

Heterologous expression in *E. coli* of the *res* and *mod* genes required the mutation of thirteen TGA codons, which codes for tryptophan in *M. bovis*, to TGG to express the full-length protein. Mod, the gene product of *mod*, could be overexpressed in *E. coli* BL21(DE3) cells and purified to homogeneity. The holoenzyme Mbo45V^0AG^/ Mbo45V^9AG^ was purified by co-expression of pRSF*mod* and pHIS*res* in *E. coli* BL21(DE3) (Figure 2A). SEC-MALS analysis of the Mod yielded a molecular mass of 131 kDa corresponding to a dimeric protein (Figure 1C). The molecular mass of both Mbo45V^0AG^ and Mbo45V^9AG^ was 225 kDa corresponding to a heterotrimeric complex of two Mod protomers and a Res subunit (Figure 1D). The oligomeric composition of Mbo45V matched with that of the prototypical EcoP15I deduced from SEC-MALS analysis and X-ray crystallography (6,7).

**Figure 2.**
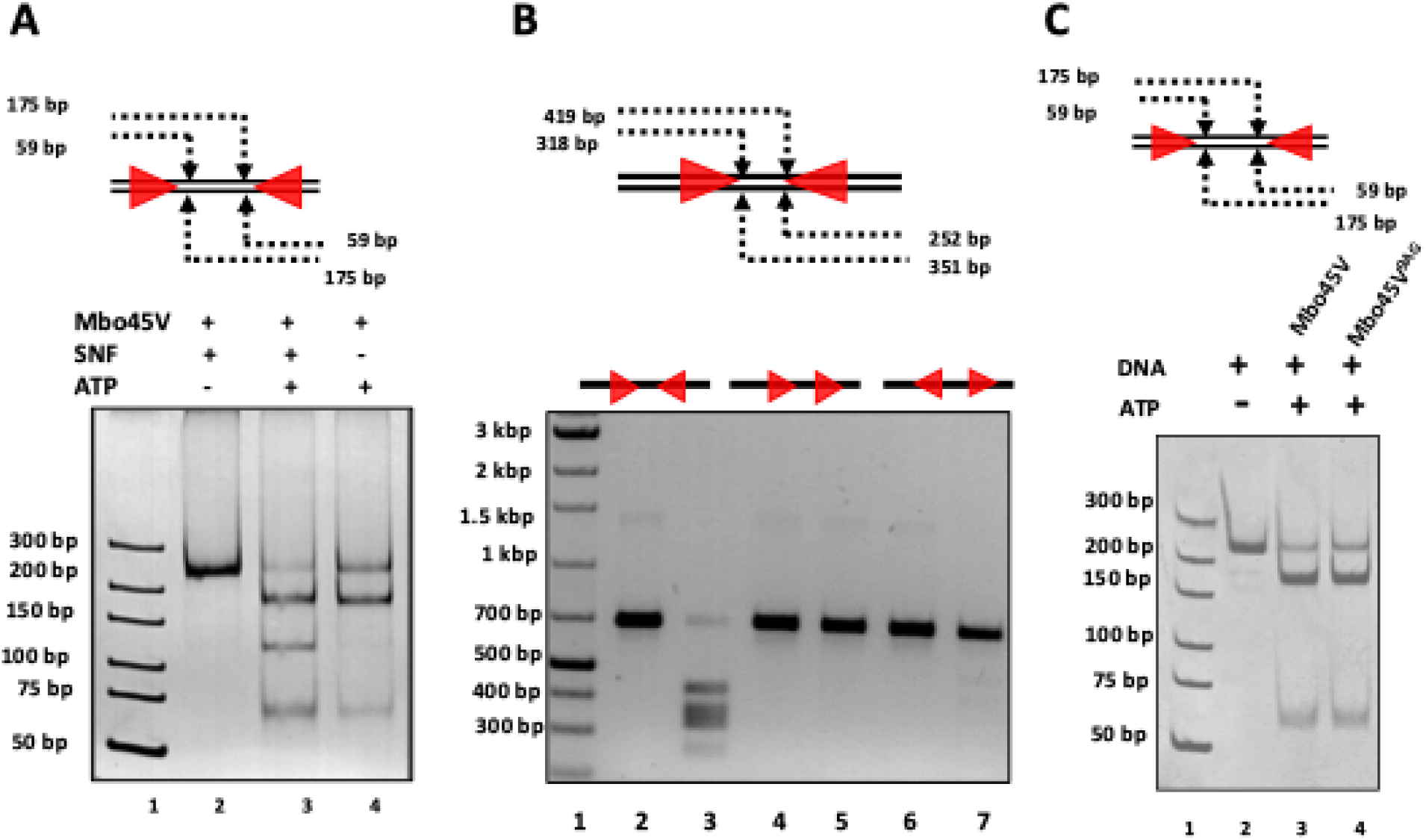
Cleavage of DNA with recognition sequence. (A) DNA containing two recognition sites (red arrowheads) in head-to-head orientation was cleaved by Mbo45V. DNA cleavage assay was carried out by incubating 300 nM Mbo45V and 50 nM of the 235 bp long DNA substrate in presence and absence of SNF. The DNA was cleaved into two fragments ∼179 bp and ∼59 bp, respectively. In presence of SNF, a band corresponding to ∼115 bp resulting from single-site cleavage is visible. (B) Site requirement for DNA cleavage by Mbo45V. DNA cleavage assay with 200 nM Mbo45V and 30 nM of a 670 bp long DNA containing two recognition sites in head-to-head orientation (Lane 3) resulted in generation of four fragments that were 481, 318, 351 and 218 bp long when resolved on a 1.5 % agarose gel. DNA substrate with recognition site in head-to-tail orientation (Lane 5) were refractory to cleavage, while the substrate with recognition sites in tail-to-tail orientation was cleaved very poorly (Lane 7). (C) DNA containing two recognition sites (red arrowheads) in head-to-head orientation was cleaved by Mbo45V contain different SSR lengths (Mbo45V^0AG^ and Mbo45V^9AG^). DNA cleavage assay was carried out by incubating 300 nM Mbo45V and 50 nM of the 235 bp long DNA. The DNA was cleaved into two fragments ∼179 bp and ∼59 bp.

### Identification of the recognition sequence of Mbo45V

We next proceeded to identify the recognition sequence of Mbo45V. For this, a set of different plasmids was treated with Mbo45V^0AG^ and checked to see if they were nucleolytically cleaved. Assuming that Mbo45V, like other known Type III RM enzymes, is an ATP-dependent endonuclease, nucleolytic cleavage of a plasmid DNA in the exclusive presence of ATP would indicate the presence of the enzyme’s recognition sites on the DNA. All the plasmids treated with Mbo45V^0AG^ were nucleolytically cleaved into several fragments in the presence of ATP, suggesting that they had multiple recognition sites of Mbo45V. Linear segments of one of these plasmids were cleaved with Mbo45V^0AG^ to screen for a segment that underwent only a single double-strand DNA break. A Type III RM enzyme’s hallmark is that it cuts DNA having at least two recognition sites in an inverted orientation. The cut occurs close to one of the recognition sites. Consequently, we assumed that the linear segment yielding only two fragments had only a pair of the Mbo45V recognition sites.

To locate the position of the recognition sites, a primer-walking assay was carried out by systematically shortening one end of the DNA while keeping the other end intact. This was repeated for both the ends. These DNA fragments were then tested for cleavage. Absence of cleavage on shortening the DNA suggested loss of a recognition site. The experiment resulted in a 218 bp DNA, which on shortening from either end prevented cleavage by Mbo45V^0AG^. Careful examination of the sequence of the DNA substrate revealed the presence of two TAATC sites oriented head-to-head. Position of the two sites was consistent with the length of DNA fragments produced on cleavage by Mbo45V^0AG^. To identify the consensus recognition sequence of Mbo45V, we generated a 235 bp DNA substrate with two TAATC sites (site A and site B in Figure 2A) in the head-to-head orientation and performed DNA cleavage assay. Each base of site B was systematically changed to the other three bases while keeping site A constant and the corresponding DNA tested for nucleolytic cleavage (Supplementary Figure 1). In total, 15 different DNA substrates were generated, and their DNA cleavage studied. Based on this study, the recognition site of Mbo45V was identified as YAATC, where Y=T/C. Best efficiency of DNA cleavage was observed in the presence of sinefungin (SNF), an analogue of the cofactor S-adenosylmethionine required for DNA methylation (Figure 2A, Supplementary Figure 2)). Previous studies with the prototypical EcoP15I had also observed efficient DNA cleavage in presence of SNF ^26^.

### Site requirement for DNA cleavage

Previous studies with EcoP1 and EcoP15I have shown that the directionality of the recognition sites is essential for DNA cleavage efficiency^27^. The recognition sites had to be in an inverted orientation for DNA cleavage, and that the head-to-tail sites were refractory to cleavage. In the context of linear DNA, recognition sites that are in head-to-head orientation are cleaved with better efficiencies than those having tail-to-tail orientation^27^. To find if the recognition sites’ relative orientations affect DNA cleavage by Mbo45V, we generated three DNA substrates with recognition sites in head-to-head, head-to-tail and tail-to-tail direction (Figure 2B). The assay revealed that the substrate with recognition sites in the head-to-head orientation was cleaved by Mbo45V^0AG^ in presence of ATP (Figure 2B, Lane 3), while that with the head-to-tail orientation was refractory to cleavage (Figure 2B, Lane 5). DNA having recognition sites in tail-to-tail direction was cleaved weakly (Figure 2B, Lane 7).

As has been reported in the case of EcoP1 and EcoP15I^9^, we also noted that Mbo45V^0AG^ could catalyze nucleolytic cleavage of the single-site substrate, but weakly, only in the presence of SNF (Figure 2A). In the absence of SNF, the single-site cleavage was unobserved. The two-site 235 bp substrate also showed single-site cleavage in the presence of SNF, resulting in an ∼115 bp DNA fragment (Figure 2A). Mbo45V^9AG^ cleaved the head-to-head two-site substrate as efficiently as Mbo45V^0AG^ (Figure 2C), indicating that the presence of SSR did not affect the enzymes nuclease activity.

To identify the location of the phosphodiester bonds hydrolyzed by Mbo45V, we performed run-off sequencing of the DNA cleavage products by Mbo45V^0AG^. The sequencing results revealed that Mbo45V cuts 26 bp 3’ to the recognition site on the YAATC strand and 28 bp 5’ to the recognition site on the GATTR strand to generate a 2 base 5’-overhang (Supplementary Figure 3). This cleavage pattern is similar to that of the prototypical Type III RM enzymes EcoP1 and EcoP15I^9^ (19).

### SNF improves target DNA binding

Studies of DNA binding of prototypical EcoP15I have established that the Type III RM enzymes have a high affinity for DNA. While DNA binding upon target recognition is mediated by the Mod dimer, the Res subunit containing the helicase and nuclease domains is expected to bind DNA non-specifically. To investigate DNA binding by Mbo45V, we performed an electrophoretic mobility shift assay (EMSA), which showed stable DNA binding that was improved with SNF (Supplementary Figure 4). Subsequently, fluorescence anisotropy studies with a 5’-FAM labelled 38 bp long DNA having target sites of Mbo45V and EcoP15I was used to measure the binding parameters (Figure 3A). The experiment yielded a dissociation constant (K_d_) of 336 nM for Mbo45V (Figure 3B). In comparison, the dissociation constant for EcoP15I was 101 nM under similar conditions (Figure 3C). In the presence of SNF, K_d_ of DNA binding by Mbo45V lowered to 207 nM (Figure 3B), while K_d_ for EcoP15I was 57 nM (Figure 3B). The analyses revealed that Mbo45V bound to its target DNA with an affinity lower than that of the prototypical EcoP15I. Also, SNF increased the binding affinity of Mbo45V, which allowed the enzyme to cleave substrate DNA better, including single site substrates.

**Figure 3.**
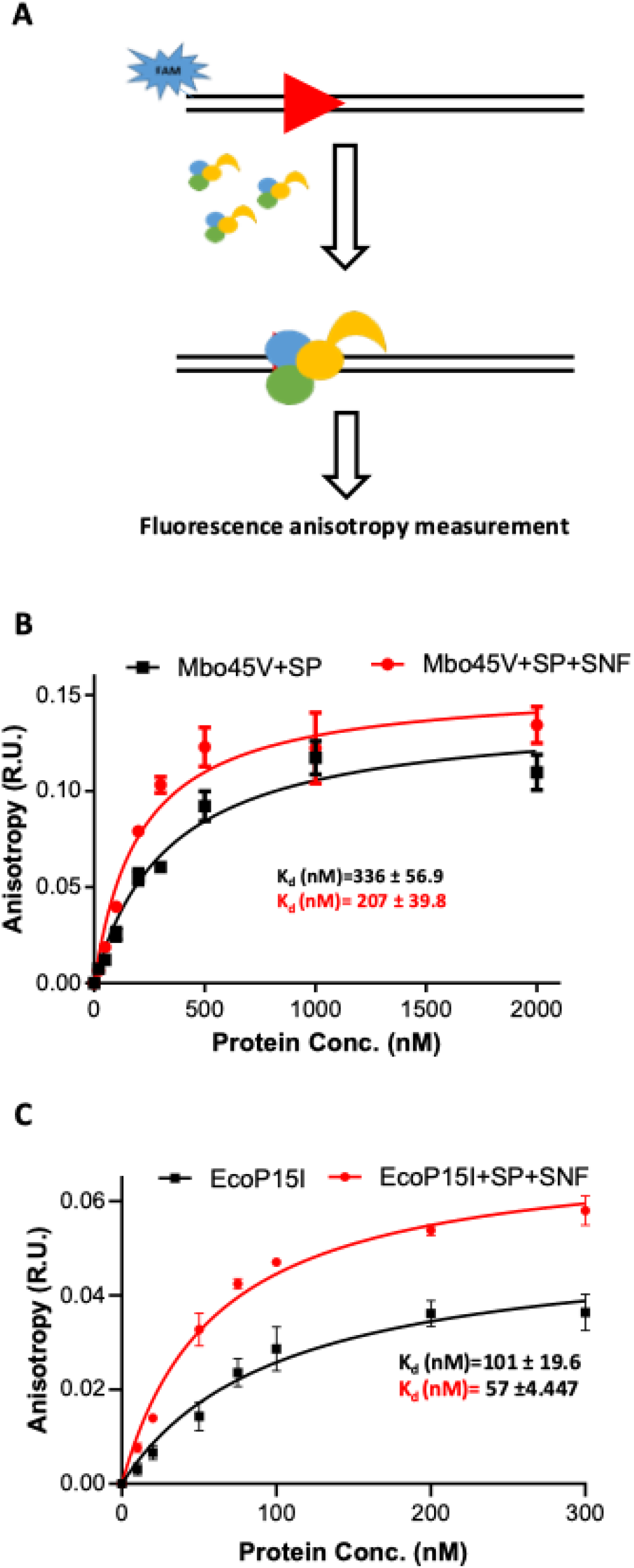
Target DNA binding of Mbo45V using fluorescence anisotropy. (A). Schematic representation of fluorescence anisotropy experiment. 50 nM of 5’ fluorescence tagged (FAM) specific DNA (SP) was incubated with variable protein concentration (10 nM – 2000 nM) then anisotropy was measured. (B) Graphical representation of fluorescence anisotropy of Mbo45V+DNA complex plotted against increasing concentration of Mbo45V with and without SNF. Data were fitted the equation for single-site specific binding. (C) Graphical representation of fluorescence anisotropy of EcoP15I+DNA complex with increasing concentration of EcoP15I with and without SNF.

### Target base for methylation by Mbo45V

The Type III RM enzymes can catalyze the methylation of their recognition site in the presence of AdoMet. Methylation of the target base is expected to prevent target recognition, and, consequently, activation of ATPase, which is dependent on target recognition. Inactivation of ATPase will inactivate DNA cleavage, which requires ATP hydrolysis. We incubated a DNA substrate having two target sites oriented head-to-head with Mbo45V^0AG^ holoenzyme in the presence of AdoMet and absence of ATP for 60 mins. After this, the DNA was purified and subjected to endonucleolytic cleavage by freshly added Mbo45V^0AG^ in the presence of ATP, and the reaction product was analyzed on a native PAGE (Figure 4A). The methylated DNA was protected from the nucleolytic activity of the freshly added Mbo45V^0AG^ (Figure 4B). Similarly, the methylation activity of Mbo45V^9AG^ were tested and found to be comparable to that of Mbo45V^0AG^ (Figure 4C).

**Figure 4.**
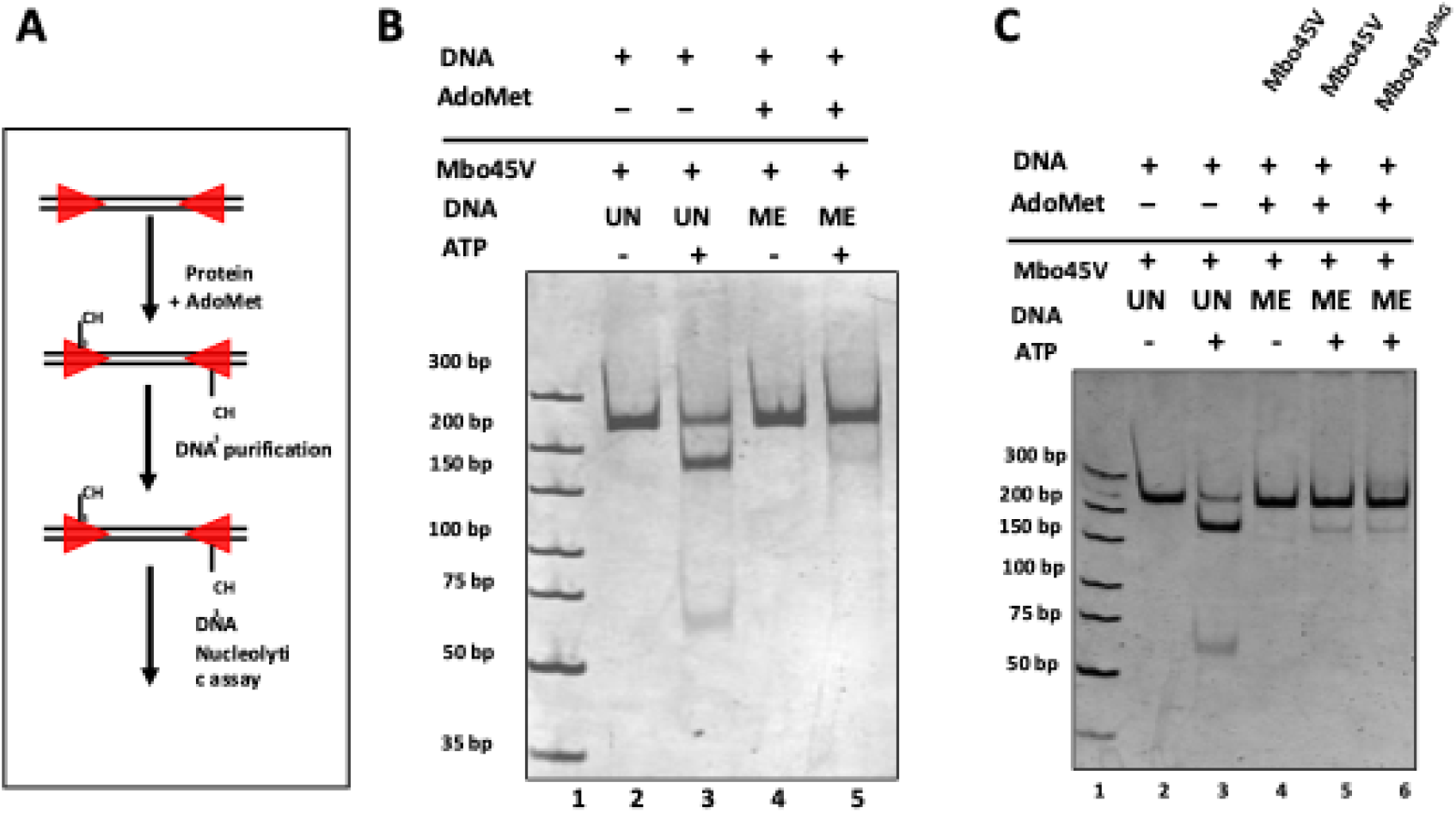
Methylation activity of Mbo45V. (A) A schematic protocol of the assay used for analyzing the methylation activity of Mbo45V. (B) Result of the assay. DNA cleavage assay was carried out with 50 nM 235 bp two-site DNA substrate that had been methylated by 300 nM of Mbo45V and 300 µM AdoMet using the protocol shown in Panel A. The methylated DNA was purified and incubated with 300 nM of fresh Mbo45V in presences of 4 mM of ATP. Methylated DNA were protected from DNA cleavage (lane 5). (C) DNA substrate were methylated with, Mbo45V^0AG^ and Mbo45V^9AG^ with AdoMet using the protocol shown in Panel A.The methylated DNA (ME) was purified and incubated with 300 nM of Mbo45V in presences of ATP to check the extent of methylation. Methylated DNA generated with Mbo45V^0AG^ and Mbo45V^9AG^ were protected from DNA cleavage.

### Kinetics of methylation reaction by Mbo45V

To further explore the MTase activity of Mbo45V, we conducted a steady-state kinetics analysis. We used a luminescence-based kit to measure the enzyme’s activity under various substrate concentrations. Methylation of target DNA by Mbo45V is a two-substrate reaction, where SAM is the methyl donor and the target DNA is the recipient. To study the kinetics of methylation, we carried out two separate experiments. In one, we varied DNA concentration (0.05-10 μM) while keeping a fixed SAM concentration of 500 μM, which is higher than the cellular concentration of SAM. The rate of methylation was plotted against DNA concentration. The Michaelis-Menten plot obtained indicated the methylation reaction rate was saturated at a DNA concentration of 3 μM or above. The kinetic parameters k_cat_ and K_m_ obtained by varying the concentration of DNA at a fixed concentration of SAM (500 μM) were 0.03 min^-1^and 0.6 μM, respectively (Figure 5A). The value of K_m_ is indicative of the affinity of the enzyme for the DNA substrate.

**Figure 5.**
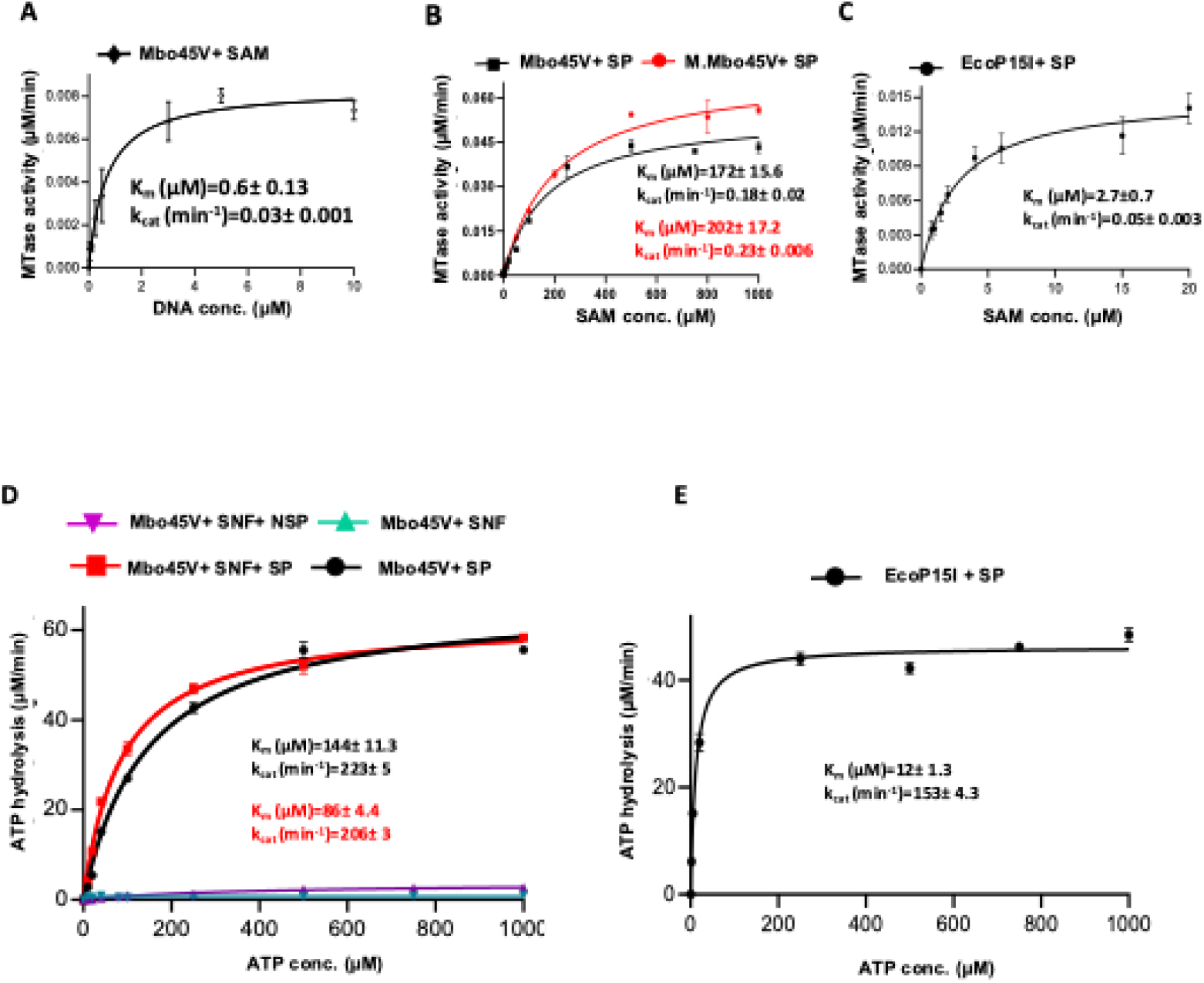
Enzyme kinetics measurement of the methylation and ATPase activity of Mbo45V. (A) Kinetics measurement of methylation activity of Mbo45V at 300 nM enzyme concentration and 500 μM of SAM. 69 bp DNA with a single target site used as substrate. Methylation activity obtained were plotted against the target DNA concentration and curve was fitted into Michaelis-Menten equation. (B) Kinetics measurement of methylation activity of Mbo45V and M.Mbo45V (in red) performed at 300 nM enzyme concentration and 3μM of 69 bp DNA with single target site. Variable SAM concentration was used to record the activity. Methylation activity obtained were plotted against SAM concentration and curve was fitted into Michaelis-Menten equation. (C) Kinetics measurement of methylation activity of EcoP15I at 300 nM enzyme concentration and 3 μM of 69 bp DNA with single target site of EcoP15I. Variable SAM concentration was used to record the activity. Methylation activity obtained were plotted against the SAM concentration and curve was fitted into Michaelis-Menten equation. (D) Kinetics measurement of ATP hydrolysis of Mbo45V at 300 nM enzyme concentration and variable concentration of ATP up to 1mM. ATP hydrolysis in presence of 1 μM of 69 bp DNA with single Mbo45V target site (SP) in black and additional SNF in red; DNA without target site (NSP) and additional SNF in green; and ATPase activity with only SNF in violet. ATPase activity obtained was plotted against ATP concentration and the curve was fitted into Michaelis-Menten equation. (E) Kinetics measurement of ATP hydrolysis of EcoP15I performed at 300 nM enzyme concentration, 1 μM of 69 bp DNA with a single target site and variable ATP concentration of ATP up to 1 mM. ATPase activity obtained were plotted against ATP concentration and the curve was fitted into Michaelis-Menten equation.

In the second experiment, methylation reaction was carried out by varying the concentration of SAM while keeping the DNA concentration constant at 3 μM. At this DNA concentration, the methylation reaction was found saturated in the first experiment. k_cat_ and K_m_ obtained from the Michaelis-Menten plot were 0.18 min^-1^ and 172.0 μM, respectively for the holoenzyme Mbo45V (Figure 5B). Here, the value of K_m_ is indicative of the affinity of the enzyme for SAM. The kinetic paramteres for the mod dimer M.Mbo45V were 0.23 min^-1^ and 202 μM (Figure 5B). Comparison between the kinetic parameters of Mbo45V and M.Mbo45V suggested that the complexation with Res did not affect the methylation activity significantly (Figure 5B). Also, kinetic studies of the methylation reaction by EcoP15I under varying SAM and constant DNA concentration, revealed that the K_m_ (SAM) of the reaction was at 2.7 μM was 64-fold lower than that for Mbo45V, while the k_cat_ was at 0.05 min^-1^ ∼3.6-fold lower (Figure 5C). This implied that Mbo45V had a very low affinity for SAM but a slightly higher methylation activity in comparison to EcoP15I. The lower affinity for SAM meant that the catalytic efficiency (k_sp_=k_cat_/K_m_) of Mbo45V for DNA methylation was much lower at 0.001 min^-1^μM^-1^ than that of EcoP15I at 0.018 min^-1^μM^-1^.

### DNA target site stimulates ATP hydrolysis

Type III RM enzymes are ATP-dependent enzymes that possess the ability to hydrolyze ATP. We used an NADH-coupled ATPase assay to investigate the ATPase activity of Mbo45V. The concentration of ATP was varied to determine the kinetic parameters for ATP hydrolysis by Mbo45V using the Michaelis-Menten plot (Figure 5D). The reactions were carried out separately in the absence of DNA with SNF, in the presence of a 69 bp specific DNA having a target site with and without SNF, and in the presence of a non-specific DNA having no target site with SNF. The plot clearly showed that Mbo45V had little ATPase activity on its own. However, addition of the cognate DNA stimulated the ATPase activity significantly. No stimulation of the ATPase activity was observed when non-cognate DNA was used. Addition of SNF improved the ATPase activity marginally only in the presence of cognate DNA. We also determined the ATPase kinetic parameters for EcoP15I in the absence of SNF (Figure 5E). While the k_cat_ and K_m_ of Mbo45V were 223 min^-1^ and 144 μM, respectively, the k_cat_ and K_m_ for the ATPase reaction by EcoP15I were 153 min^-1^ and 12 μM respectively. This implied that Mbo45V had a much lower affinity for ATP than EcoP15I. Accordingly, the catalytic efficiency, k_sp_, of the ATPase activity of Mbo45V was approximately 8-fold lower than that of EcoP15I.

### Methylation of DNA by Mbo45V inhibits its nucleolytic activity

We next proceeded to study the effect of methylation on the ATPase and nuclease activities of Mbo45V, and to find which of the two reactions, methylation or ATPase, is more prominent when both SAM and ATP are available in the reaction mix. Mbo45V can catalyze either methylation or cleavage of the DNA substrate depending on the availability of the cofactor SAM or ATP, respectively. To find which of the two reactions would be prominent if both the cofactors were present simultaneously, we performed an assay in which the DNA substrate was incubated with Mbo45V^0AG^, 4 mM ATP and 200 µM AdoMet, simultaneously. Methylation of the target base is expected to prevent target recognition, and, consequently, prevent activation of ATPase, which is dependent on target recognition, and, consequently affect ATP-dependent DNA cleavage. After incubation for 60 minutes, the DNA was analyzed on a native PAGE. We found that most of the fractions were protected from cleavage by methylation (Supplementary Figure 5). This suggested that methylation was kinetically faster than nucleolytic cleavage.

Next, we carried out a set of cleavage reactions at a constant ATP concentration, while increasing the SAM concentration from 0 to 300 μM stepwise. We observed that an increase in the SAM concentration reduced the cleavage activity. As the concentration of SAM approached 300 μM, cleavage of DNA by Mbo45V diminished considerably (Figure 6A). In contrast to Mbo45V, EcoP15I showed almost no cleavage activity at a much lower SAM concentration of 0.7 μM (Figure 6B).

**Figure 6.**
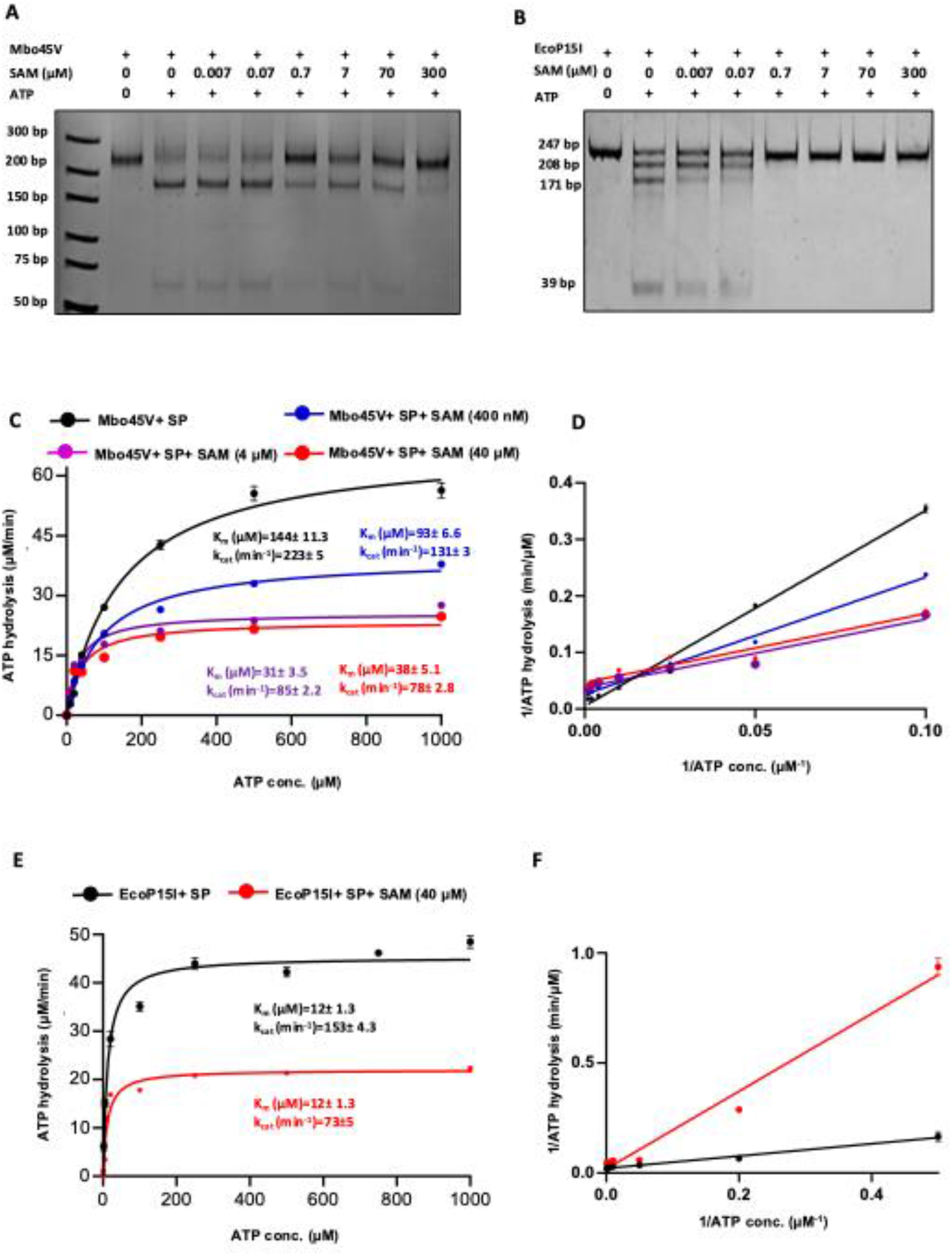
Analysis of competing activity of Mbo45V. (A) Competition assay between methylation and nuclease activities of Mbo45V in which 50 nM 235 bp two-site DNA substrate was incubated with 300 nM of the enzyme in the presence of increasing SAM concentration (0.007 μM – 300 μM) and ATP (1 mM). In the presence of 300 μM SAM concentration, DNA cleavage was diminished, showed the DNA substrate was methylated before the nucleolytic cleavage at higher concentration. (B) Competition assay between methylation and nuclease activities of EcoP15I in which 50 nM 224 bp two-site DNA substrate was incubated with 300 nM of the enzyme in the presence of increasing SAM concentration (0.007 μM – 300 μM) and ATP (1mM). In presence of 0.7 μM SAM concentration or above, DNA cleavage was diminished. (C) Kinetics measurement of ATP hydrolysis of 300 nM Mbo45V enzyme with 1 μM of 69 bp DNA with target site and variable concentration of ATP up to 1 mM was performed with different SAM concentration. The ATPase activity plotted against ATP concentration and fitted to Michaelis-Menten equation. The enzyme’s kinetic graph showed inhibition with increasing SAM concentration. The curve without SAM is in black, Michaelis-Menten curve at 400 nM SAM concentration represented in blue, at 4 μM SAM concentration represented in magenta and at 40 μM SAM concentration represented in red. (D) Observed activity data of Mbo45V was plotted for Lineweaver-Burk plot. (E) Kinetics measurement of ATP hydrolysis of 300 nM EcoP15I enzyme with 1 μM of 69 bp DNA with target site and variable concentration of ATP up to 1mM was performed with two different SAM concentrations. The ATPase activity plotted against the ATP concentration and fitted to Michaelis-Menten equation. The enzymes kinetic graph at different SAM concentration showed inhibition upon increasing SAM concentration. The curve without SAM is in black, and Michaelis-Menten curve at 40 μM SAM concentration represented in red. (F) Observed activity data of EcoP15I was plotted for Lineweaver-Burk plot.

We also performed ATPase assay of Mbo45V with varying concentrations of SAM to find how methylation of the DNA substrate affected the kinetics of the ATPase activity. We found that the ATPase activity of Mbo45V decreased with increasing concentration of SAM and the kinetics followed the Michaelis-Menten reaction mechanism. The parameters K_m_ and k_cat_ decreased with increasing SAM concentration in a stepwise manner from 0 μM to 40 μM (Figure 6C). The decrease in k_cat_ as well as K_m_ indicated a more complex mechanism of mixed inhibition of ATPase due to the methylation of the DNA substrate in the presence of SAM. In the corresponding Lineweaver-Bruk plot (figure 6D), the lines intersected in the first quadrant, suggesting that the product of the methylation reaction, i.e. the methylated DNA, bound to enzyme with high affinity and prevented ATP hydrolysis^28^. Fluorescence anisotropy measurement yielded a binding constant of 400 nM (Supplementary Figure 6). As a control, we carried out the kinetic study of the ATPase in presence of the SAM analogue SNF. We found that k_cat_ changed very little, while the K_m_ for the ATPase reaction decreased significantly in comparison to that for the ATPase reaction carried out in the presence of SAM (Figure 5D).

When we performed the same kinetic study with EcoP15I, we observed that the k_cat_ of the ATPase decreased when the concentration of SAM was changed in a stepwise manner, while the K_m_ for ATP remained constant (Figure 6E, 6F). This suggested that, in the case of EcoP15I, the inhibition of the ATPase reaction caused by DNA methylation reaction appeared non-competitive. As the methodology used by us to measure the kinetics of methylation involved detection of production of ATP, we could not measure the kinetics of methylation in presence of varying concentration of ATP.

## Discussion

The biochemical characterization of Mbo45V reported by us identifies it as a bona fide Type III RM enzyme. Although Mbo45V shares low sequence identity with the prototypical Type III RM enzymes EcoP15I and EcoP1, it exhibits very similar biochemical properties. Like the prototypical EcoP1 and EcoP15I, Mbo45V is an oligomer composed of two Mod and one Res polypeptides. The enzyme cleaves DNA containing at least two recognition sequences, 5’-YAATC-3’, arranged in an inverted orientation, with substrates harboring sites in a head-to-head orientation being cleaved significantly more efficiently than those with sites in a tail-to-tail orientation.

Consistent with the prototypes, cleavage occurs ∼26–28 bp downstream of the recognition sequence, resulting in a 2-nucleotide 5′ overhang. The enzyme also displays weak nucleolytic activity on single-site substrates. Like EcoP15I^26,29^, Mbo45V shows improved target DNA binding and cleavage activities in the presence of SNF. Apart from the similarities, our biochemical studies reveal that the kinetic parameters of the enzymatic activities of Mbo45V from the host-adapted obligate parasite *M. bovis* are quite different from those of EcoP15I or EcoP1, prophage encoded enzymes from *E. coli*, which is a facultative anaerobe that inhabits a wide range of environments.

In comparison to EcoP15I or other N^6^-adenine DNA MTases that have DNA dissociation constant (K_d_) in the range of 0.1 to 60 nM^30^ (E.coli/T4 DAM, CcrM, CamA), Mbo45V exhibited a significantly high K_d_ of 336 nM in the absence of SNF and 207 nM in presence of SNF for cognate DNA. The Michaelis constant K_m_ measured for the methylation reaction carried out by varying DNA concentration while keeping the SAM concentration constant was about 3-fold higher at 600 nM. K_m_ being higher than K_d_ could reflect rate limiting step/s following DNA binding, such as base flipping and protein conformation changes, which occurs when EcoP15I binds to its cognate DNA to form a catalytically competent state^31^, and is expected to occur when Mbo45V binds to its cognate DNA too.

The methylation reaction catalyzed by Mbo45V was kinetically distinct from that catalyzed by EcoP15I when measured at varying concentrations of SAM and a constant concentration of DNA substrate. While Mbo45V showed a 4-fold higher *k*_cat_ compared to EcoP15I, its K_*m*_ (SAM) of approximately 173 μM was nearly 50-fold higher than that of EcoP15I, implying a considerably weaker affinity for SAM under steady state conditions. This implies that Mbo45V will efficiently methylate DNA only when SAM concentration is as high as hundreds of micromolar. In contrast, the low *K*_m_ (SAM) value of ∼2 μM of EcoP15I indicates that the enzyme will function effectively as an MTase even at low cellular concentrations of SAM.

The lower affinity of Mbo45V for SAM likely reflects adaptation to the metabolically active and nutrient-rich intracellular host environment in which *M. bovis* exists as an obligate parasite, as well as to the relatively low to moderate bacteriophage exposure. The approximate intracellular concentration of SAM in mycoplasma (*M. pneumoniae*) is 0.004% of cell mass (∼100 μM)^32^. Consequently, Mbo45V can tolerate higher K_m_ values due to a relatively stable cofactor concentrations. In contrast, SAM concentration in *E. coli* are known to vary from ∼30 μM to 230 μM and can occasionally reach as high as 10 mM within the organism depending on its growth phase and the fluctuating environmental conditions it is living in^33^. MTases from *E. coli*, including EcoP15I, display much higher affinity for SAM^30^, (E.coli/T4 DAM^34^, CcrM^35^, CamA^36^). This links the methylation activity to cellular physiology through cofactor availability. The kinetic parameters of the ATPase reaction catalyzed by Mbo45V followed a trend similar to that observed for the methylation reaction. In comparison to EcoP15I, Mbo45V possesses a significantly higher *K*_m_ and a comparable *k*_cat_ value, implying higher ATPase activity under conditions where cellular ATP concentrations are high in millimolar range.

Results from the cleavage competition assays of Mbo45V and EcoP15I reported here explain these kinetic differences, as DNA cleavage by Mbo45V is favored at most of the SAM concentrations tested and tapers out only at higher concentrations of SAM. In contrast, EcoP15I-mediated DNA cleavage is blocked even at 0.7 μM SAM, a consequence of the low K_m_ of its MTase activity. Assuming methylation of the substrate DNA to be inhibitory, from a Lineweaver-Bruk plot, found that the inhibition of the ATPase upon DNA methylation was mixed. A mixed inhibition with intersection of the lines in the first quadrant suggested that the methylated DNA remained bound to the enzyme complex and diminished ATP hydrolysis and consequently DNA cleavage. Contrastingly, inhibition was non-competitive in the case of EcoP15I, where again the inhibitory methylated DNA remains bound to the enzyme.

In summary, the results presented and discussed through this study demonstrate that the methylation and ATPase activities of Mbo45V are kinetically distinct from those of EcoP15I. This likely reflects the environmental niche occupied by the obligate parasite *M. bovis* in comparison to the facultative anaerobe *E. coli* that inhabits diverse environments - from nutrient-rich to nutrient-poor and from phage-free to phage-rich conditions.

## Notes

### Competing Interest Statement

The authors have declared no competing interest.

