## Supplementary material for "Biochemical and kinetic properties of a Type III restriction-modification enzyme Mbo45V from the host-adapted pathogen *Mycoplasma bovis*": NA

| | T | A | G | C |
|---|---|---|---|---|
| T | + | - | - | + |
| A | - | + | - | - |
| A | - | + | - | - |
| T | + | - | - | - |
| C | - | - | - | + |

**Supplementary Figure 1.** Different DNA substrates generated for DNA nuclease assay when recognition site at each base was changed to other three bases. DNA cleavage happened (marked by a tick) only when recognition site was (YAATC, where Y = T/C).

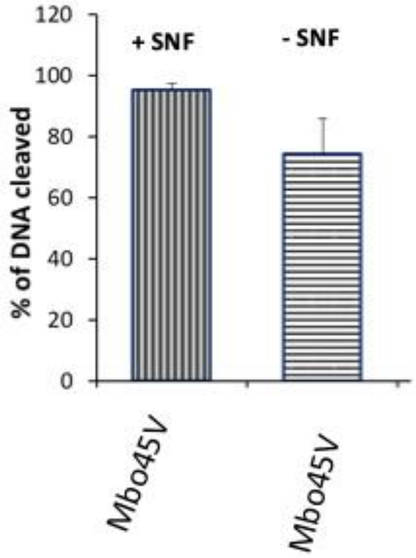


**Supplementary Figure 2.** Quantification of DNA cleavage shown in the representative gel in 2A. DNA bands were quantified using Image J software. The plot represents data from three independent experiments.

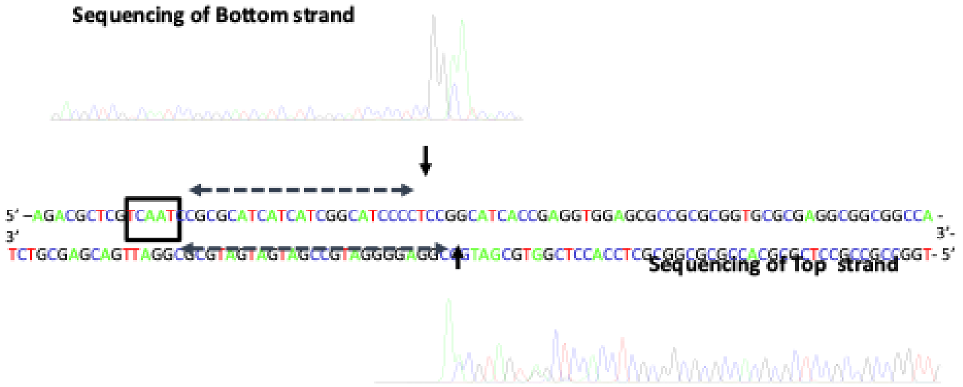


**Supplementary Figure 3.** Run-off sequencing to determine the cleavage loci of Mbo45V. The recognition sequence is marked by a rectangular box. The arrows in the top and bottom strands indicate the position of the two nicks. Mbo45V nicks the top strand 26 bp downstream of the recognition sequence and bottom strand is nicked 28 bp downstream of the recognition sequence.

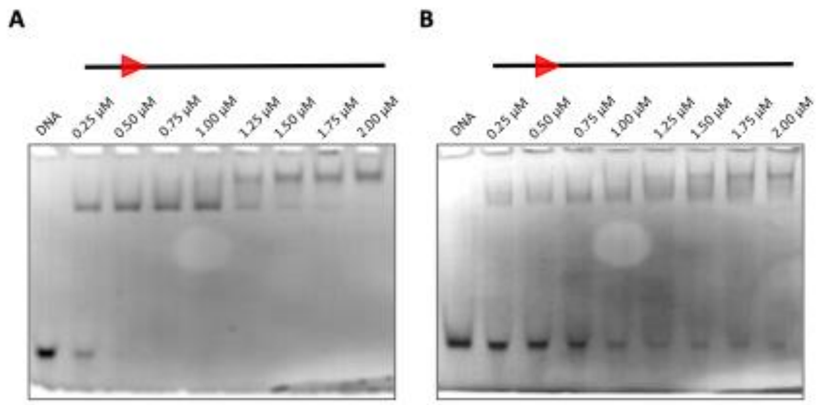


**Supplementary Figure 4.** DNA binding of SMbo45V to a 30 bp specific DNA in the presence and absence of SNF analyzed using EMSA.

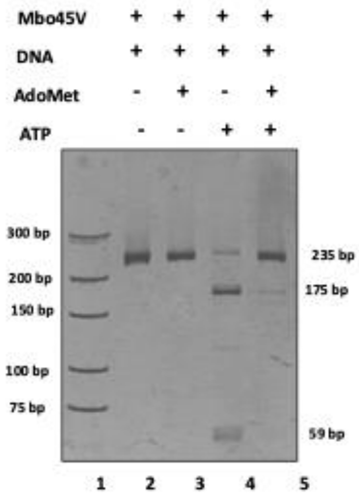


**Supplementary Figure 5.** Competition assay between methylation and nuclease activities of Mbo45V in which 50 nM 235 bp two-site DNA substrate was incubated with 300 nM of the enzyme was incubated in presence of 200 μM AdoMet and 4 mM ATP. The presence of AdoMet diminished DNA cleavage, suggesting that the DNA substrate was methylated before the nucleolytic cleavage could occur.

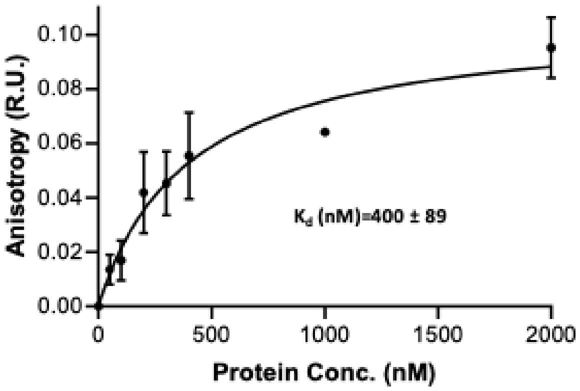


**Supplementary Figure 6.** Graphical representation of fluorescence anisotropy of Mbo45V+ methylated DNA complex plotted against increasing concentration of Mbo45V.
